# A Chemical-genetics and Nanoparticle Enabled Approach for *in vivo* Protein Kinase Analysis

**DOI:** 10.1101/2020.05.13.094573

**Authors:** Fengqian Chen, Qi Liu, Terrell Hilliard, Tingzeng Wang, Hongjun Liang, Weimin Gao, Leaf Huang, Degeng Wang

## Abstract

The human kinome contains >500 protein kinases, and regulates up to 30% of the proteome. Kinase study is currently hindered by a lack of *in vivo* analysis approaches due to two factors: our inability to distinguish the kinase reaction of interest from those of other kinases in live cells and the cell impermeability of the ATP analogs. Herein, we tackled this issue by combining the widely used chemical genetic method developed by Dr. Kevan Shokat and colleagues with nanoparticle-mediated intracellular delivery of the ATP analog. The critical AKT1 protein kinase, which has been successfully studied with the method, was used as our initial prototype. Briefly, enlargement of the ATP binding pocket, by mutating the gate-keeper Methionine residue to a Glycine, prompted the mutant AKT1 to preferentially use the bulky ATP analog N^6^-Benzyl-ATP-γ-S (A*TPγS) and, thus, differentiating AKT1-catalyzed and other phosphorylation events. The lipid/calcium/phosphate (LCP) nanoparticle was used for efficient intracellular delivery of A*TPγS, overcoming the cell impermeability issue. The mutant, but not wild-type, AKT1 used the delivered A*TPγS for autophosphorylation and phosphorylating its substrates in live cells. Thus, an *in vivo* protein kinase analysis method has been developed. The strategy should be widely applicable to other protein kinases.

## 1. Introduction

Protein kinases function by cleaving the γ-phosphate group off ATP and covalently adding the group onto serine, threonine or tyrosine residues of their substrate proteins, up or down-regulating their activities in vital cellular processes such as cell cycles, growth, migration and apoptosis (1,2). Very often, the substrates themselves are protein kinases. Through this phosphorylation process, the protein kinases form many signaling cascades, *e*.*g*., the MAP kinase cascades (3). The kinases and their cascades constitute the backbone of the cellular signaling network, which bridges extracellular and intracellular environment and interconnects subcellular compartments/domains. Many diseases exhibit aberrancies in protein kinases, *i*.*e*., over-activation of disease promoting kinases and/or over-repression of disease suppressive ones (4,5). Thus, the treatments often target these kinases. For instance, the type II diabetic treatment targets the AMPK protein kinase (6), and cancer treatments aim to correct the aberrancies of oncogenic and/or tumor-suppressive protein kinases (3,5).

However, kinase study continues to be hindered by a major challenge – insufficient *in vivo* analysis capability. The tremendous efforts over the last two decades have led to fruitful *in vivo* activity monitoring for some protein kinases (7-10). Such methods generally work via the expression of a fusion protein composed of a sensing domain, a reporter domain and often a linker region. The sensing domain contains the phosphorylation site of the kinase of interest, phosphorylation of which results in readily quantifiable effects on the activity of the reporter domain. Examples of the effect includes activation of FRET, and changes of fluorescence (intensity, wavelength or subcellular locations), activities. However, these strategies have significant drawbacks. First, when homologous kinases share the same substrates, as is often the case, these methods are powerless in differentiating them. Second, they monitor the equilibrium of opposite effects of kinase and phosphatase activities on the sensing domain, instead of solely the kinase activity. Third, a protein kinase often has a large number of substrates with varying affinity to it. The activity on the site chosen for the sensing domain might not reflect faithfully the activities on all of the substrates. Fourth, the methods have no power in identifying new substrates. In a word, our ability for *in vivo* analysis is far from satisfactory yet.

Currently, kinase studies still rely heavily on one *in vitro* strategy – incubation of ATP with labeled γ-phosphate group, un-phosphorylated peptide substrate and the kinase in the reaction buffer followed by detecting/quantifying substrate tagging by the labeled γ-phosphate group from ATP. The kinase can be generated through recombinant expression followed by purification and, in case one wishes to quantify an endogenous kinase activity, isolation from cellular extracts. However, it is very much questionable whether the *in vitro* substrate tagging reliably reflects the *in vivo* phosphorylation events and their magnitudes.

In order to establish *in vivo* kinase analyses in intact live cells, we need to meet two requirements. First, it is necessary to be able to distinguish the kinase reaction of interest from those of the other protein kinases, as the human kinome contains more than 500 protein kinases (2). Second, we need to develop a method to generate sufficient intracellular abundance of the tagged ATP analog. The analog by itself is not cell permeable, but is required as a probe to detect and/or monitor the kinase reactions, *i*.*e*., transfer of the γ-phosphate group from ATP onto the substrates.

Fortunately, the well established chemical-genetic strategy for identifying kinase-substrates relationships developed by Dr. Kevan Shokat’s lab is well suited to accomplish the first requirement. The Shokat method uses site-directed mutagenesis to enlarge a kinase’s ATP binding pocket. While not affecting kinase activity, the mutation specifically allows bulky synthetic ATP analogs, such as N^6^-Benzyl-ATP (A*TP), to fit into the enlarged ATP binding pocket (11-16). Moreover, A*TP was further modified to create A*TP-γ-S by replacing an O with a S in the γ-phosphate group. The mutant protein kinase preferentially use A*TP-gamma-S in its kinase reaction, transferring the thiophosphate tag from A*TP-γ-S onto its substrate proteins. This tag can then be alkylated by a chemical called *p*-nitrobenzylmesylate (PNBM) to generate a thiophosphate-ester for specific antibody recognition or affinity purification followed by LC-MS analysis (17-19). That is, this method combines ATP-binding pocket enlargement and the bulky ATP analog A*TP-gamma-S to distinguish the kinase reaction of interest from those of the other protein kinases, enabling identification of the substrates of specific kinases. However, due to the cell impermeability of the A*TP-γ-S molecule, this method have been executed *in vitro* with cell lysate in test tubes. In order for the thiophosphate-tagging of the substrates to occur in live cells, we need to accomplish the second requirement, *i*.*e*., a reliable method for efficient intracellular A*TP-γ-S delivery.

As for this second requirement, a tool also seems exist already. The lipid/calcium/phosphate (LCP) nanoparticle is well suited for this task. It consists of a calcium phosphate (CaP) core, which carries the to-be-delivered chemicals, and a lipid bilayer, which encapsulates the CaP core and enables cellular uptake of the nanoparticle through endocytosis. This nanoparticle has been used for efficient intracellular delivery of Gemcitabine Triphosphate, a nucleotide analog used as a chemotherapy agent (20,21). The A*TP-γ-S molecule is chemically similar to Gemcitabine Triphosphate. Thus, we became interested in using the LCP nanoparticle as a vehicle for intracellular A*TP-γ-S delivery and, in turn, to enable intracellular thiophosphate-tagging of protein kinase substrates via the Shokat chemical genetic approach; that is, achievement of *in vivo* protein kinase analysis via a combination of the Shokat chemical genetic method and the nanoparticle delivery of the A*TP-γ-S molecule.

To accomplish this goal, we choosed the serine/threonine protein kinase AKT1 (also known as PKB) as the initial prototype. Firstly, AKT1 is functionally and therapeutically important (22). It is part of the critical phosphoinositide 3-kinase (PI3K) signaling module (23-27). The non-receptor tyrosine kinase ACK1/TNK2 can also phosphorylate AKT1, resulting in PI3K-independent activation (28). AKT1 phosphorylates and regulates a wide range of metabolic and/or regulatory substrate proteins such as IKKalpha, GSK3beta, and FOXO (29,30). Thus, it participates in multiple vital cellular processes, including cell survival and cell growth. As the Akt signaling process aberrancy occurs in multiple human diseases such as diabetes and cancer, AKT1 is a significant therapeutic target for multiple human diseases (22,24,29,31-36). Secondly, perhaps due to the functional importance, AKT1 is one of the earliest protein kinases studied successfully with the Shokat chemical genetic approach, enhancing the likelihood of success of this study (37-39). Moreover, in addition to phosphorylating its numerous substrates, AKT1 undergoes auto-phosphorylation (40), further facilitating this initial study.

Thus, following the Shokat method, the AKT1 ATP binding pocket was enlarged by mutating the gate-keeper Methionine residue into a Glycine. We also used the LCP nanoparticle for efficient intracellular delivery of the A*TPγS molecule, overcoming the cell impermeability issue. We demonstrated that the mutant, but not the wild type, AKT1 was able to use the delivered A*TPγS for autophosphorylation as well as phosphorylating other protein substrates in live cells. In a word, an *in vivo* AKT1 kinase analysis method has been developed, to the best of our knowledge, for the first time. The strategy uses only basic commodity reagents and equipments, and should be widely applicable to other protein kinases.

## 2. Materials and methods

### 2.1 Materials

Alexa-Fluor-647-ATP was obtained from Thermo Fisher Scientific (Massachusetts, USA), and N^6^-Benzyladenosine- 5’- (3-thiotriphosphate) (A*TPγS) was obtained from BIOLOG Life Science Institute (Bremen, Germany). IKK alpha monoclonal antibody was obtained from Thermo Fisher Scientific (Massachusetts, USA), anti-thiophosphate ester antibody was obtained from Abcam (Cambridge, UK), and AccuRuler RGB protein ladder and BP-Fectin mammalian cell transfection reagent were obtained from BioPioneer (California, USA). Dioleoyl phosphatydic acid (DOPA), 1,2-dimyristoyl-3-trimethylammonium-propane chloride salt (DOTAP), and 1,2-distearoyl-sn-glycero-3-phosphoethanolamine-N-[amino(polyethylene glycol)-2000] ammonium salt (DSPE-PEG_2000_) were obtained from Avanti Polar Lipids, Inc. (Alabama, USA). DSPE-PEG-anisamide (AEAA) was synthesized in Dr. Leaf Huang’s lab as described previously (41). Other chemicals were purchased from Sigma-Aldrich, Inc. (Missouri, USA).

### 2.2 Cell culture

HCT116 human colon cancer cells, originally obtained from Dr. Michael Brattain’s lab (42,43), were cultured in the McCoy’s 5A medium (Invitrogen, USA) supplemented with 10% fetal bovine serum, 100 U/mL penicillin, and 100 mg/mL streptomycin (Invitrogen, USA). Cells were cultivated at 37 °C and 5 % CO_2_ in an incubator.

### 2.3 Preparation of A*TPγS-loaded LCP nanoparticles

Briefly, 100 µL of 10 mM specified ATP analog solution (A*TPγS or Alexa-Fluor-647-ATP) in 12.5 mM Na_2_HPO_4_ solution and 100 µL of 2.5 M CaCl_2_ solution were dispersed in cyclohexane/Igepal solution (70/30, v/v), respectively, to form an oil phase. After 15 min of stirring, 100 µL of 25 mM DOPA was added into the oil phase for 15 min, then 40 mL of 100 % ethanol was added into the oil phase. Later, the mixture solution was centrifuged at 10,000 g for 20 min to pellet the LCP particle core out of the supernatant solution. After washing with 50 mL ethanol twice, 100 µL of 10 mg/mL cholesterol, 100 µL of 25 mg/mL DOTAP, 100 µL of 25 mg/ml DSPE-PEG, and 10 µL of 25 mg/mL DSPE-PEG-AEAA were added into the LCP core solution. After evaporating under N_2_, the residual lipids were suspended in PBS to produce the layer of LCP nanoparticle. After being sonicated for 10 min, the LCP solution was dialyzed in PBS.

### 2.4 Characterization of A*TPγS-loaded LCP nanoparticles

The LCP nanoparticle particle sizes was determined by Malvern DLS (Royston, UK). The concentration of A*TPγS was determined by an HPLC spectrophotometer (Shimadzu Corp., Japan). Transmission electron microscope (TEM) photos of A*TPγS-loaded LCP nanoparticles were observed under JEOL 100CX II TEM (Tokyo, Japan).

### 2.5 Intracellular uptake behaviors

Intracellular uptake behaviors were observed with a confocal microscopy. Briefly, the HCT116 cells were plated into confocal dishes (1 × 10^5^ cells per well) and incubated for 24 hours. Then, the cells were incubated with Alex-Fluor-647-ATP-loaded LCP nanoparticles for 4 hours with a concentration of 5 µg/mL, and were nucleus-labeled with Hoechst 33342 dye (Thermo Fisher Scientific) for 15 min. The images were taken with a Zeiss 880 confocal microscope (Germany).

### 2.6 Plasmid constructs

Wild type AKT1 plasmid was purchased from Addgene (Addgene plasmid pcDNA3-myr-HA-AKT1 1036). Point mutation as in references (37) of the ATP-binding pocket gatekeeper amino acid (M227G) were introduced by GENEWIZ (New Jersey, USA), with a modified Stratgene QuikChange® site-directed mutagenesis method.

### 2.7 Expression plasmid transfection

The HCT116 cells were plated into 100 mm^2^ dishes and transfected at approximately 90% confluence with the specified plasmid (wild type pcDNA3-myr-HA-AKT1 plasmid or the mutant pcDNA3-myr-HA-AKT1 plasmid) using BP-Fectin mammalian cell transfection reagent (BioPioneer, USA) in accordance with the manufacturer’s instructions. One untransected dish was used as a negative control. After 72 h, cells were washed twice with cold PBS. Transfection efficacy was confirmed by western-blot measurement of the expression of the HA-tagged fusion proteins with the HA tag monoclonal antibody (Thermo Fisher Scientific, USA).

### 2.8 Intracellular A*TPγS delivery and thio-phosphorylation assays

Control and transfected HCT116 cells were incubated with A*TPγS-loaded LCP nanoparticles for 4 hours. The cells were washed by cold PBS twice and then lysed with RIPA buffer (which contains protease and phosphatase inhibitors) for 30 minutes at 4 °C. The cell lystaes were treated with 2.5 mM PNBM at room temperature for 1 hour. Then, the PNBM treated cell lysates were analyzed with a western blot assay to measure protein thio-phosphorylation with the anti-thiophosphate ester rabbit monoclonal antibodies (Abcam, USA).

### 2.9 Immunoprecipitation assays

The PNBM-treated cell lysates were pre-cleaned by an off-target antibody. Then the cleaned cell lysates were incubated with 10 µL anti-thiophosphate ester rabbit monoclonal antibody (Abcam, USA) at 4 °C for 12 h. 100 µL protein G beads (Thermo Fisher Scientific, USA) were mixed under rotary agitation at 4 °C for 4 h, washed twice by wash buffer, and the beads were collected. Then, proteins were captured in the beads and released by elution buffer. The elution buffer was collected and stored at −80 °C.

### 2.10 Western blotting assay

First, total protein concentrations of the cell lysates were determined by a BCA protein assay kit (Thermo, USA). Protein extracts were separated by SDS–PAGE, then transferred onto a nitrocellulose membrane (NC membrane). After blocking the NC membrane with 5 % non-fat milk for 1 h at room temperature, it was incubated with the specified first antibody (the HA tag monoclonal antibody, the anti-thiophosphate ester antibody or the IKK alpha monoclonal antibody) at 1:1000 dilution at 4 °C overnight. Membranes were washed five times with TBST solution, incubated with the corresponding secondary antibody (1:3000 dilution) for 1.5 h at normal room temperature, and washed five times with TBST solution. Protein bands were visualized using an enhanced chemiluminescence system according to the manufacturer’s instructions (Thermo Fisher Scientific, USA).

### 2.11 PhosphoSitePlus dataset analysis

In order to perform human kinome-wide analysis, the PhosphoSitePlus dataset was downloaded from the database website (www.phosphosite.org). For this study, all non-human protein kinases and substrates were excluded. Substrate count was then calculated for each kinase with the remaining data.

## 3. Results

### 3.1 Characterization of A*TPγS-loaded LCP Nanoparticles

As mentioned earlier, the LCP nanoparticle is a formulation that has been used for efficient intracellular delivery of the nucleotide analog Gemcitabine Triphosphate as a chemotherapy agent. Herein, we tested it for intracellular A*TPγS delivery to overcome the cell impermeability issue. Figure 1 shows a schematic illustration of the preparation procedure of A*TPγS-loaded LCP nanoparticles. A*TPγS is first loaded into the CaP core of the LCP nanoparticles, which is then coated with a DOPA layer. DOPA functions to prevent the aggregation of the CaP core because of the strong interaction between the CaP core and the phosphate head of DOPA. The cationic 1,2-dioleoyl-3-trimethylammonium-propane (DOTAP) lipid is then added as an outer layer onto the DOPA layer (21). Additionally, an outside layer of neutral dioleoylphosphatidylcholine (DOPC) with DSPE-PEG with a tethered targeting ligand anisamide (AA) is added. These layers keep the nanoparticles stable in the hydrophilic tissue culture medium and also enhance cellular uptake of the nanoparticles. Upon nanoparticle cellular internalization, the CaP core is rapidly dissolved in the endosome due to its low pH, leading to increased endosome osmotic pressure. Consequently, the endosome swells and ruptures, releasing the loaded A*TPγS into the cytoplasm. This quick process also avoids lysosomal degradation after endocytosis.

**Figure 1.**
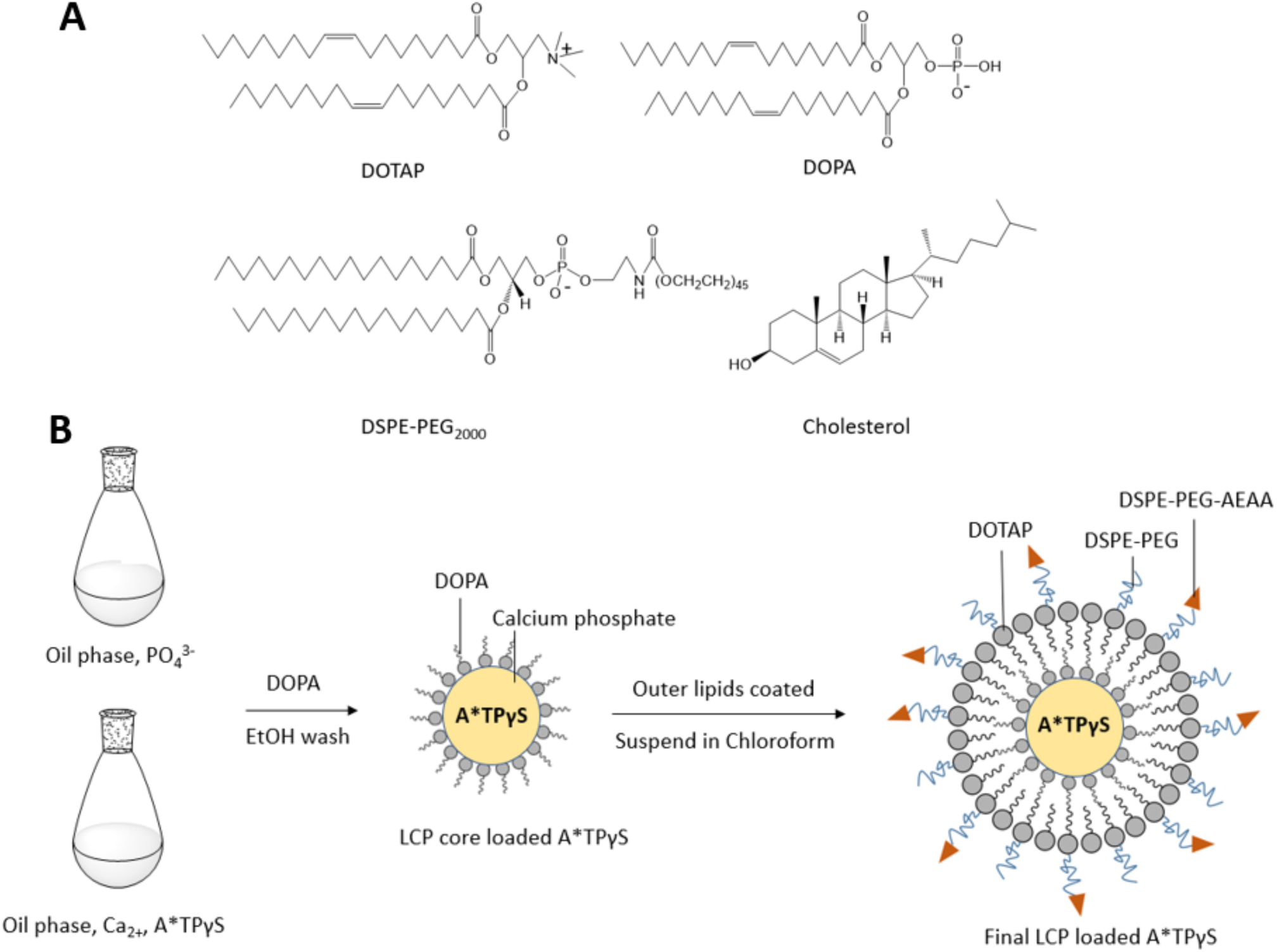
LCP nanoparticle and its synthesis. **A:** Chemical structures of DOTAP, DOPA, DSPE-PEG and cholesterol. **B:** Schematic illustration of the synthesis procedure of A*TPγS-loaded LCP nanoparticles.

Figure 2 summarized the results of our characterization of synthesized nanoparticles. The loaded LCP nanoparticle sizes are exhibited in Figure 2A in a histogram. The sizes are also summarized in Figure 2B along with those of unloaded nanoparticles. On average, A*TPγS-loaded LCP nanoparticles were 83.7±8.1 nm and, as expected, about 11% larger than unloaded nanoparticles, which has a size of 75.1±5.2 nm. A transmission electron microscope (TEM) image is also shown in Figure 2C, illustrating the monodispersed nanoparticles with spherical shapes. The final concentration of A*TPγS in 1 mL of produced nanoparticles was 0.5 mM. Thus, about 50% of the starting A*TPγS (100 µL of 10 mM solution) was loaded into the nanoparticles.

**Figure 2.**
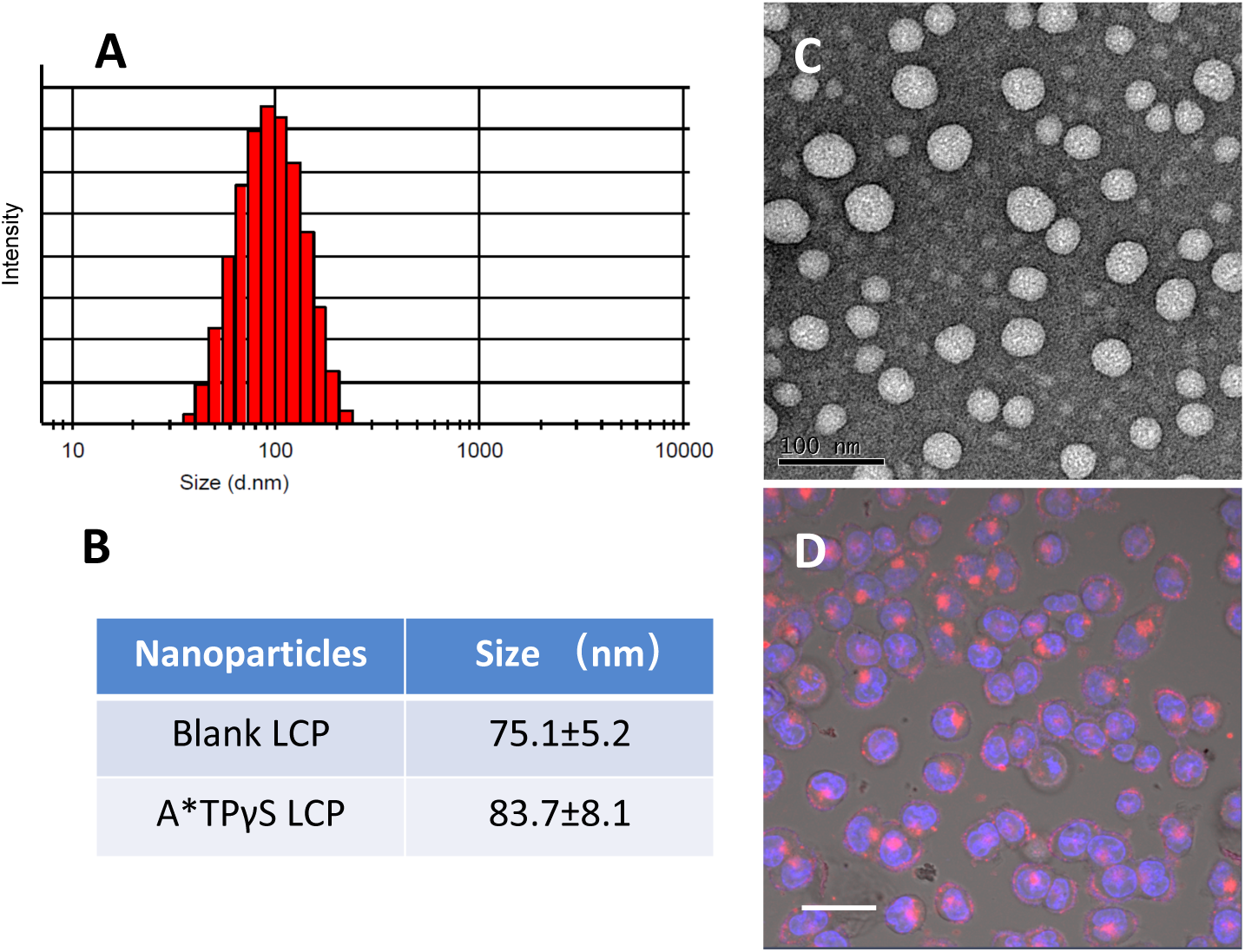
Characterization of the LCP nanoparticles. **A:** Histogram of nanoparticle size. **B:** Average sizes of blank LCP nanoparticles and A*TPγS-loaded LCP nanoparticles. **C:** TEM images of A*TPγS-loaded LCP nanoparticles. The experiment was repeated 6 times. A representative result is shown. The scale bar is 100 nm. **D:** Internalization of Alexa-Fluor-647-ATP-loaded LCP nanoparticles in HCT116 cells. The experiment was repeated 4 times. A representative result is shown. The scale bar is 20.0 µm.

We also monitored intracellular uptake of the LCP nanoparticle by using the fluorescent ATP analog Alexa-Fluor-647-ATP. As described in Materials and Methods, this ATP analog was loaded into the LCP nanoparticles. The loaded nanoparticle was added into the tissue culture medium and incubated with the HCT116 cells for 4 hours for intracellular uptake. The cells were then nucleus-stained and observed under a confocal microscope. The result is shown in Figure 2D. Strong Alexa-Fluor-647 red fluorescence activities were spotted inside the HCT116 cells, demonstrating the cellular internalization of loaded LCP nanoparticles. Thus, we were convinced that the LCP nanoparticles could be used as a vehicle for intracellular A*TPγS delivery.

### 3.2 Generation and expression of mutant AKT1 kinase

Meanwhile, we generated other components of the Shokat chemical genetic method, in order to combine it with nanoparticle-mediated intracellular A*TPγS delivery to enable *in vivo* protein kinase substrate detection.

First, as brifly mentioned earlier, the AKT1 protein kinase was chosen as our initial prototype to develop an *in vivo* method for identifying protein kinase-substrates relationship due to feasibility and functional importance. The AKT1 mutant with enlarged ATP binding pocket has been shown to bind to the bulky ATP analogue A*TP. The mutant retained the protein kinase activity and exhibited the same ectopic expression level as the wild type AKT1 kinase (37,38). It was also reported by Dr. Shokat’s group that ATPγS (note: not A*TPγS) can be utilized by wild type AKT1 to tag its substrate GSK3β *in vitro* (44). In a word, these evidences support AKT1 as the most feasible choice of initial prototype. Moreover, Figure 3 illustrated quantitatively the functional importance of AKT1 in terms of the number of experimentally determined AKT1 subtrates. We downloaded the PhosphoSitePlus database (www.phosphosite.org) dataset and counted the number of unique experimentally verified substrates for each human protein kinases, revealing that human AKT1 has 215 unique substrates (45). As many other cell biology parameters (46-49), the substrate count (K) of the human kinome follows the so-called scale-free distribution, *i*.*e*., P_(k)_ ∝ K^−α^ or log_2_(P_(k)_) ∝ -α*log_2_(K), with P_(K)_ as the portion of protein kinases with K unique substrates and α as a positive constant. This is illustrated in the linear relationship in a log-log plot (log_2_(P_(k)_ vs. log_2_(K)) in Figure 3. A small number of protein kinases have extra-ordinarily high substrate counts, while the majority of the kinases have only a small number of substrates. As shown by the red * symbol in Figure 3, AKT1 is one of the high-substrate-count kinases; it was ranked, as shown in the insert table in Figure 3, at the 7^th^ among all human protein kinases. Therefore, AKT1 is not only the most feasible, but also a functionally important, choice for our study.

**Figure 3.**
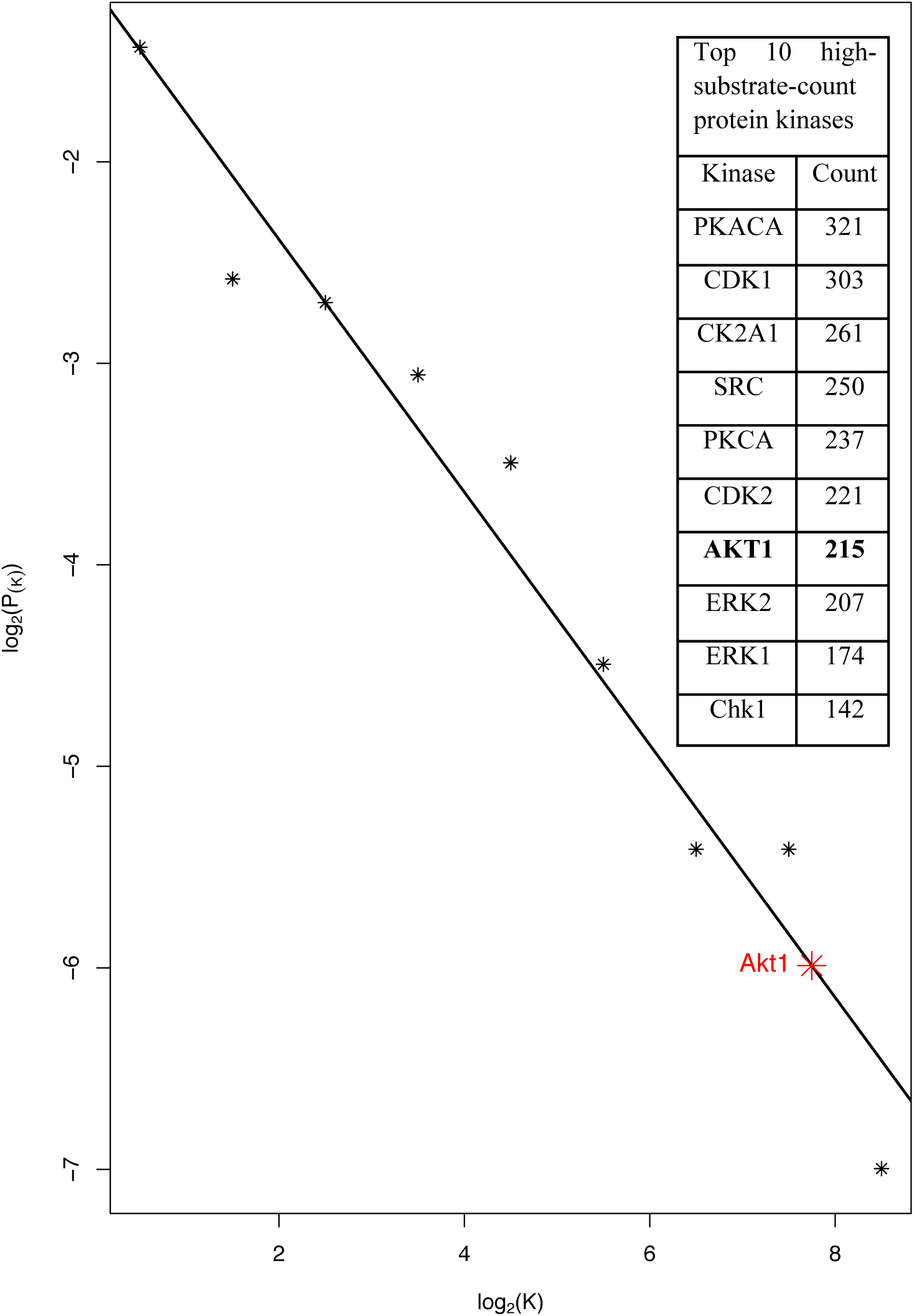
High substrate count of AKT1. A log-log plot of the portion of the kinases with K substrates (P_(K)_) and the substrate count (K) depicts the scale-free distribution of K, *i*.*e*., a linear relationship between log_2_(P_(K)_) and log_2_(K). The red * symbol and text illustrate the high Akt1 substrate count. The insert table lists the top 10 high-substrate-count kinases, with AKT1 in bolded text.

Thus, we acquired the pCDNA3-Myr-HA-AKT1 expression plasmid from AddGene. The vector expresses the AKT1 protein fused to the HA and the Myr tags. The HA-tag enables detection of the expressed AKT1 protein with the HA-tag antibody, facilitating transfection analysis. The Myr tag renders the expressed protein myristoylated, membrane anchored and, thus, contitutively active. We applied site-directed mutagenesis to mutate the gatekeeper Methionine codon into a Glycine codon in AKT1 ATP binding pocket (37). The two vectors made it possible for us to test whether the bulky A*TPγS molecule is too large for AKT1 and other wild type protein kinases, but fits into the enlarged ATP binding pocket of mutant AKT1 and leads to specific thiophosphorylation of AKT1 substrates.

Second, to test thio-phosphate-tagging of substrate proteins in live cells, a host cell line was needed. The human HCT116 cell line were chosen. This cell line has an activating point mutation in the PI3KCA gene (50,51), resulting in constitutive PI3K kinase activity and thus elevated activation of the expressed AKT1 fusion proteins, further facilitating *in vivo* detection of thio-phosphate-tagging of AKT1 substrates.

Having established these components, we next tested the expression of both wildtype and mutant AKT1 proteins upon transfection of the respective expression plasmids into the HCT116 cells and subsequent treatments. 72 hours after the tranfection, the cells were treated with the A*TPγS-loaded nanoparticle for 4 hours. The cells were then lysed. The cell lysates were treated with the PNBM alkylation to convert the thiophosphate tag into thiophosphate-ester tag, and then aliquoted. We then performed Western blot analysis with one aliquote set of the cell lysates and the HA-tag antibody to meaure the ectopic expression of wildtype and mutant AKT1 proteins. As shown in Figure 4A, both wild type and mutant AKT1 expression vectors drove strong expression of the HA-tagged proteins in the HCT116 cells upon transfection, while the control/untransfected HCT116 cells exhibited no detectable HA-tagged AKT1 expression. The A*TPγS-loaded nanoparticle and the PNBM treatments did not affect the expression significantly. The other aliquots of the same cell lysates were used, as described below, in subsequent downstream analyses

**Figure 4.**
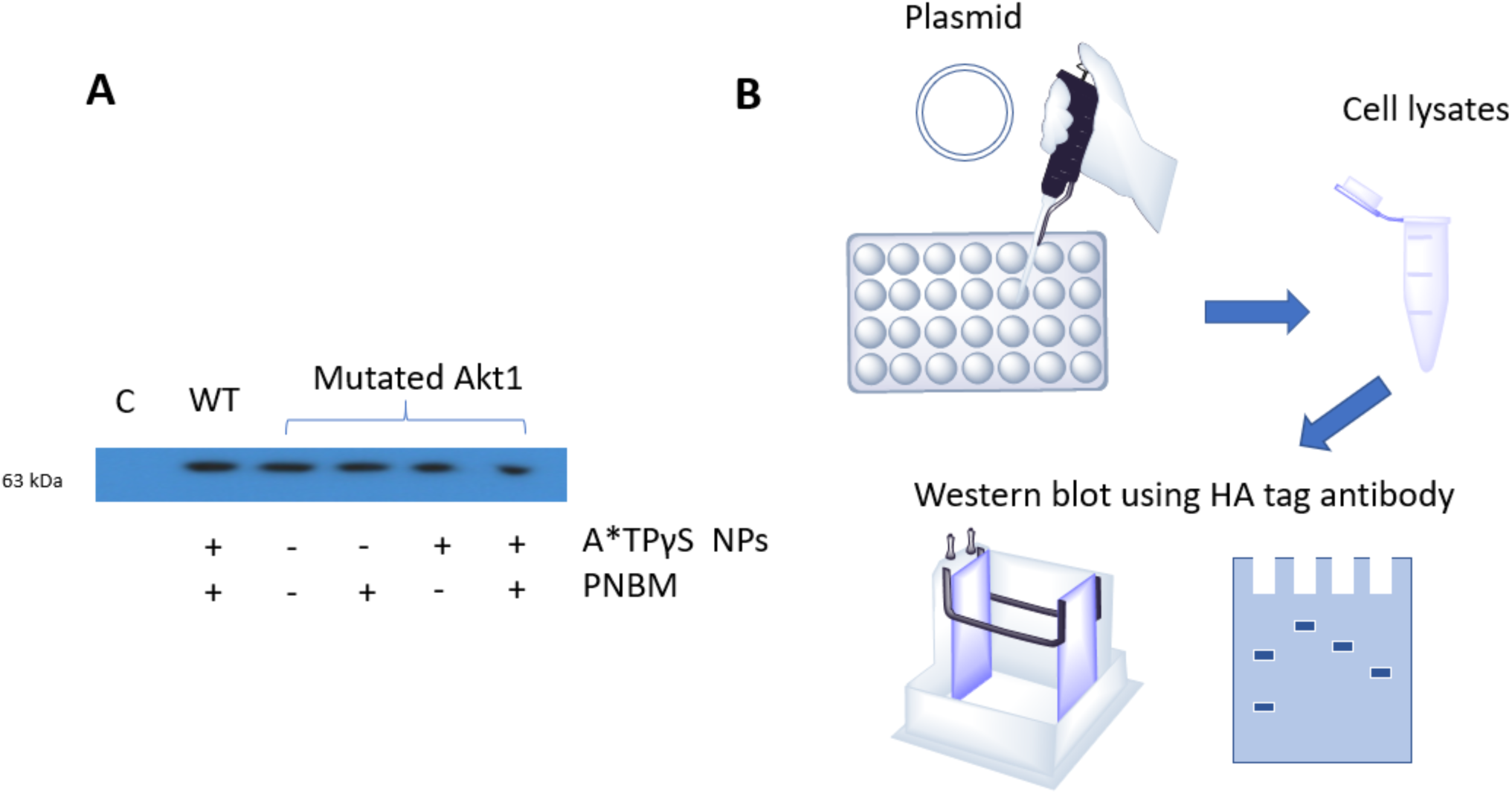
Expression of AKT1 proteins from transfected plasmid vectors. **A:** HA-tag Western blot analysis of control/untransfected HCT116 cells (C) and HCT116 cells transfected with either the wild type (WT) or the mutant AKT1 expression plasmids (Mutant Akt1). The cells were treated with the A*TPγS-loaded LCP nanoparticles and then lysed. The cell lysates were alkylated by PNBM and subjected to Western blot analysis with the anti-HA-tag antibody. The experiment was repeated more than 3 times. A representative result is shown. **B:** Transfection and western blot analyses workflow.

### 3.3 Specific *in vivo* utilization of A*TPγS by mutant AKT1

Next, we attempted to determine whether the mutant AKT1 protein can use the nanoparticle-delivered A*TPγS in its phosphorylation reaction, *i*.*e*., transferring the γ-thiophosphate group onto AKT1 substrates (Figure 5). For this purpose, we tested potential AKT1 self-tagging through its autophosphorylation activity as well as tagging of another well-known AKT1 substrate, the IKKalpha protein (30). Since alkylation by PNBM should have converted the thiophosphate tag into the thiophosphate ester tag (52), the anti-thiophosphate ester monoclonal antibody should be able to recognize the tagged proteins. Thus, another aliquot set of the treated cell lysates was used for the tests. The lysates of the transfected cells were subjected to immunoprecipitation with the thiophosphate ester tag antibody. The immunoprecipitants, along with the un-immunoprecipitated lysate of control/untransfected cells, are then examined by Western blot analyses with HA-tag or IKKalpha antibodies; the un-immunoprecipitated control/untransfected cell lysate served as negative and positive controls for the HA-tag and the IKKalpha antibodies, respectively. The results are shown in Figure 6. The HA-tag antibody was used in this analysis to detect self-tagging of the Myr-HA-AKT1 fusion proteins due to AKT1 auto-phosphorylation. As shown in Figure 6A, the fusion protein was detected in the immunoprecipitant of the cells transfected with the mutant AKT1 expression vector. Both nanoparticle treatment of the cells and PNBM alkylation of the cell lysate were required; the bands disappeared when either treatment was not performed. However, the fusion protein was not detected in the immunoprecipitants of the cells transfected with wild type AKT1 expression vector, even though the cells were treated with A*TPγS-loaded LCP nanoparticle and the cell lysate was also treated with PNBM. Not surprisingly, the control/untransfected cell lysate gave no band due to a lack of expression of the HA-tagged fusion protein. The same was observed for the IKKalpha protein (Fig. 6B). IKKalpha was detected with the IKKalpha antibody in the immunoprecipitants of the cells transfected with the mutant AKT1 expression vector, but not in that of the cells transfected with wild type AKT1 expression vector. As expected, the antibody was able to detect endogenous IKKalpha protein in the un-immunoprecipitated control/untransfected HCT116 cell lysate. Thus, the mutant AKT1 protein was able to use, exclusively, the nanoparticle delivered intracellular A*TPγS for *in vivo* thiophosphate-tagging of its substrates in live HCT116 cells.

**Figure 5.**
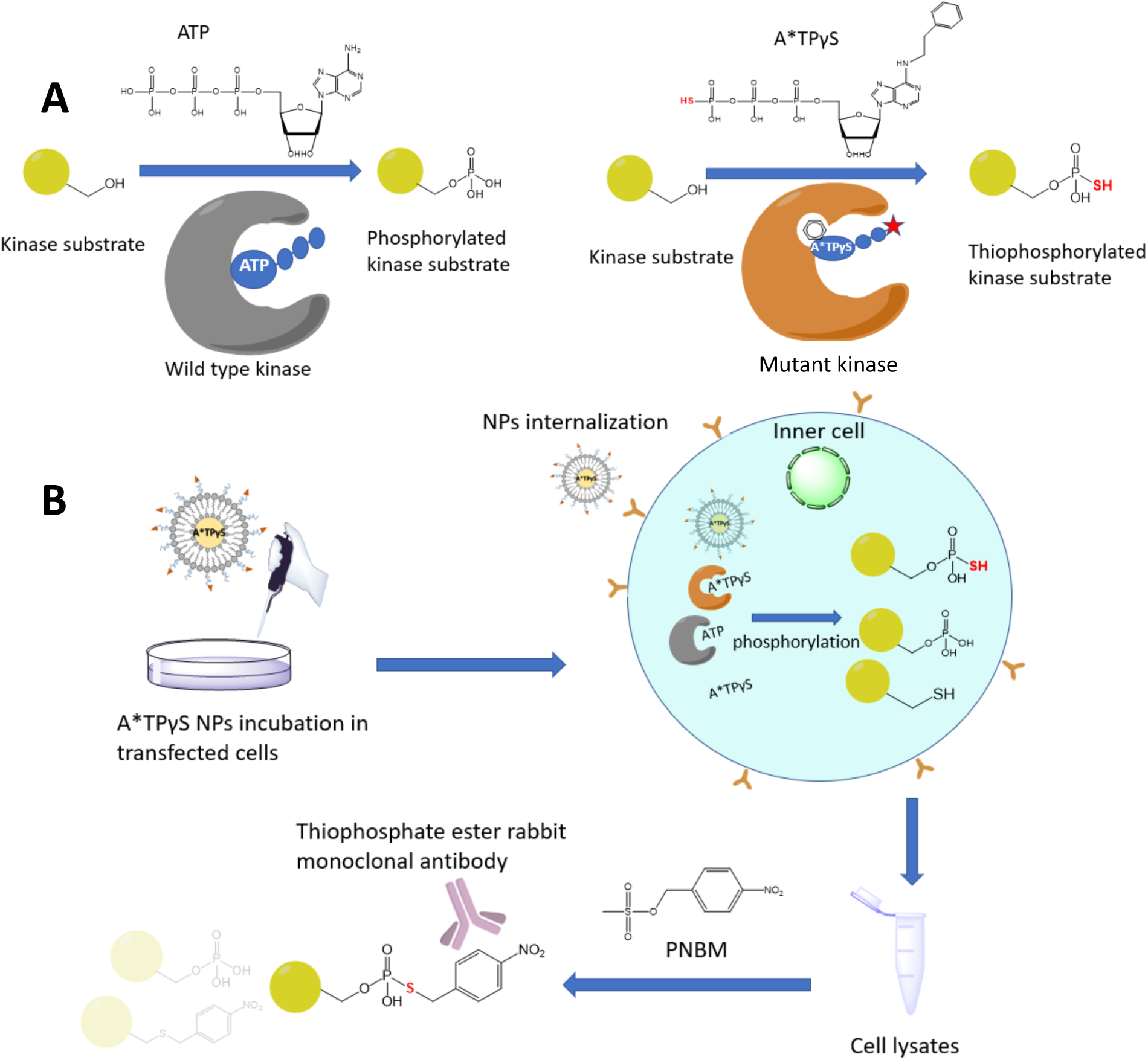
*In vivo* thiophosphate-tagging of kinase substrates and its detection strategy. **A:** Wild type kinase utilizes normal ATP to phosphorylate its substrates. In contrast, the mutated kinase, with enlarged ATP binding pocket, accepts bulky A*TPγS and was able to transfer the thiophosphate group onto the substrates. **B:** Workflow for thiophosphate-labelling of kinase substrates.

**Figure 6.**
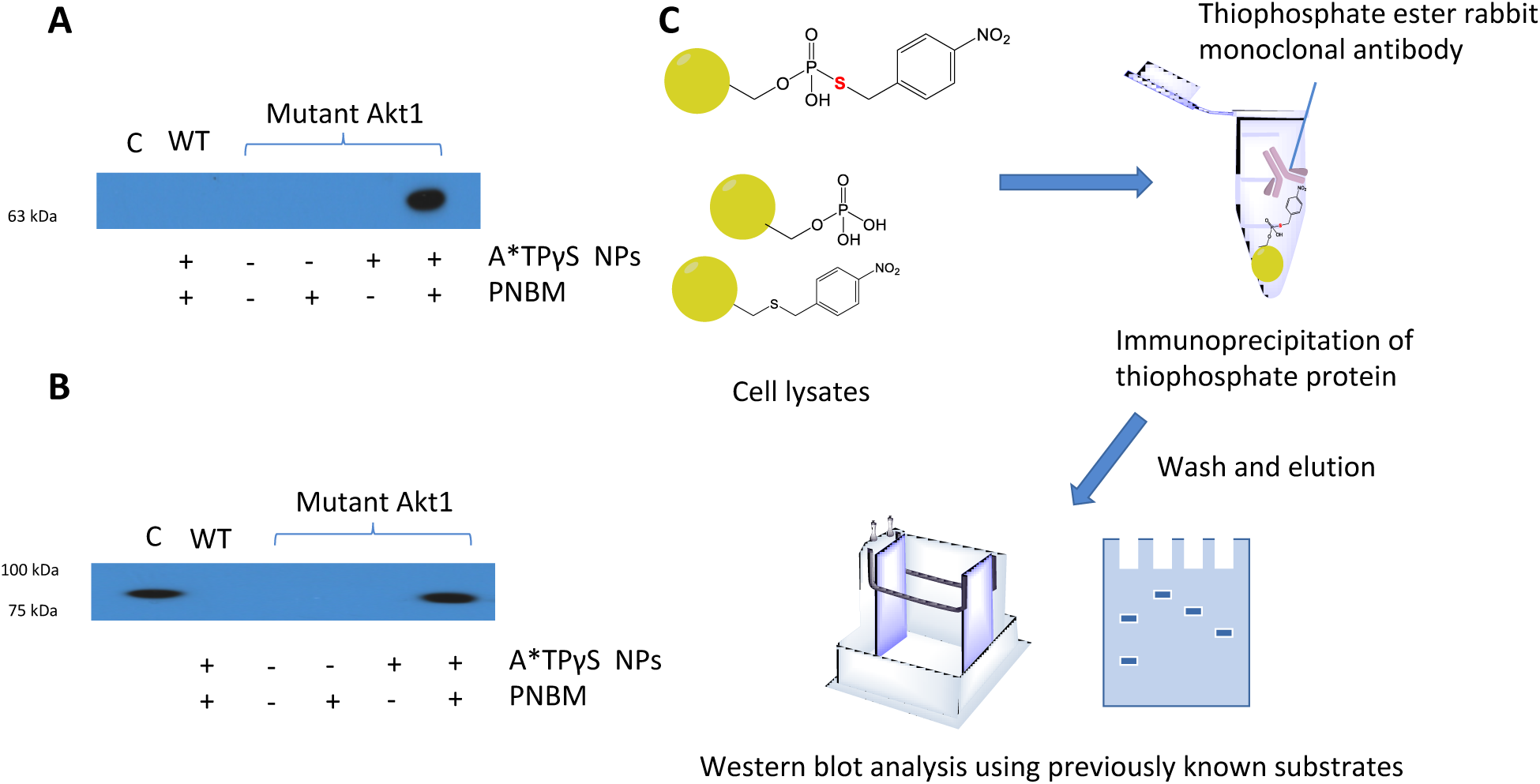
Analyses of *in vivo* AKT1 auto-thiophosphorylation and thiophosphorylation of AKT1 substrate IKKα. **A** and **B**: The transfected HCT116 cell lysates were subjected to anti-thiophosphate-ester antibody immunoprecipitation. This was followed by HA-tag (**A**) or IKKα (**B**) Western blot analyses of the control/untransfected HCT116 transfected cell lysate (C) as well as the immunoprecipitants of the lysates of HCT116 cells transfected with either wildtype (WT) or mutant AKT1 (Mutant Akt1) expression plasmids. The same cell lysates as in figure 4 were used. Please note that the control/untransfected HCT116 cell lysate was directly used without immunoprecipitation. The experiments was repeated 3 times. A representative result is shown. **C:** Schematic illustration of the experimental workflow of immunoprecipitation of thiophosphated proteins followed by Western blot analyses.

AKT1 is known to phosphorylate a large number of substrates. As discussed earlier, a search of the PhosphoSitePlus database (www.phosphosite.org) revealed that human AKT1 has 215 unique human proteins as its experimentally verified substrates and was the 7^th^ human kinase with most substrates (45). Thus, we next tried to determine whether the mutant AKT1 protein would use the delivered A*TPγS to tag many other proteins in the HCT116 cells. First, we tried Western blot with the thiophosphate-ester antibody to detect the presence of tagged proteins in another aliquot set of the cell lysates, which had been used in Figures 4 and 6. As shown in Figure 7, only the cells that were transfected with the mutant AKT1 expression vector and treated with A*TPγS-loaded nanoparticles and whose cell lysates were PNBM-alkylated exhibited bands in the Western blot analysis (Figure 7). Multiple prominent bands were detected, ranging from 25 to180 kDa. This is not surprising, given that AKT1 is known to phosphorylate a large number of substrates. As expected, wild type AKT1 was not able to use the bulky A*TPγS, although the cell lysate was also treated with PNBM. Once again, for cells transfected with the mutated AKT1 expression vector, both nanoparticle treatment of the cells and PNBM alkylation of the cell lysate were required; the bands dispeared when either treatment was not performed. Second, we performed immunoprecipitation of the PNBM alkylated lysates of cells transfected with expression vectors and treated with the nanoparticle. The immunoprecipitants were subjected to SDS-PAGE electrophoresis. The gel was then silver blue comassie stained, in order to compare the protein abundance in the two immunoprecipitants. As expected, the thiophosphate ester antibody was able to immnoprecipitate at high quantities a large number of proteins from the lysate of mutant-AKT1-expressing cells, but not from that of wild-type-AKT1-expressing cells (Figure 8). Taken together, the results of the two experiments suggests that LCP-nanoparticle delivered intracellular A*TPγS can be used exclusively by the mutant AKT1 protein to thiophosphate-tag its substrates in live cells. That is, by combining nanoparticle-mediated intracellular A*TPγS delivery and the Shokat chemical genetic method, an *in vivo* approach for kinase-substrate relationship determination has been established.

**Figure 7.**
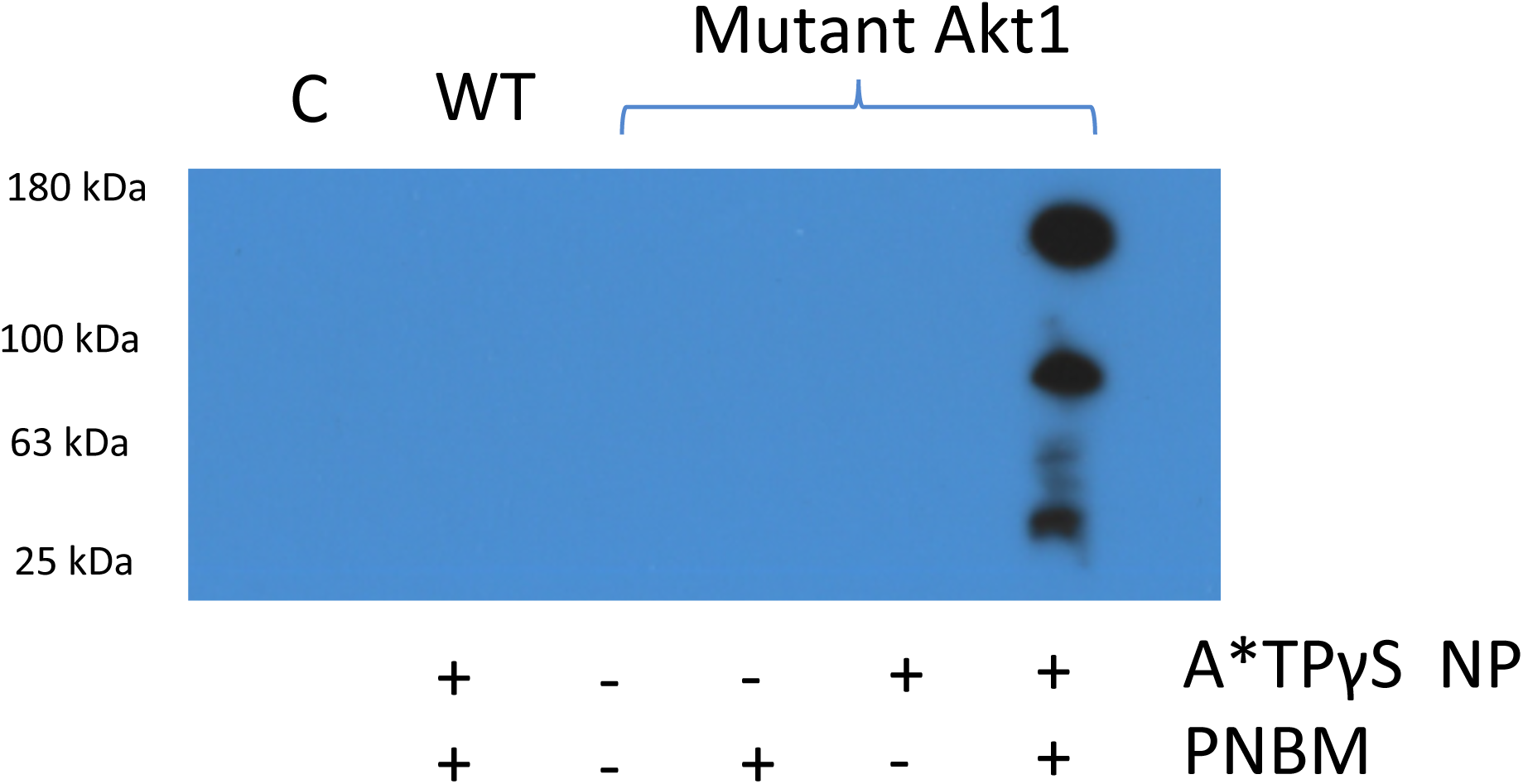
Western blot analysis of thiophosphorylated proteins. The same cell lysates as in figures 4 and 6 were used. The PNBM-alkylated cell lysates were subjected to Western blot analysis with the anti-thiophosphate-ester antibody. The experiment was repeated 3 times. A representative result is shown.

**Figure 8.**
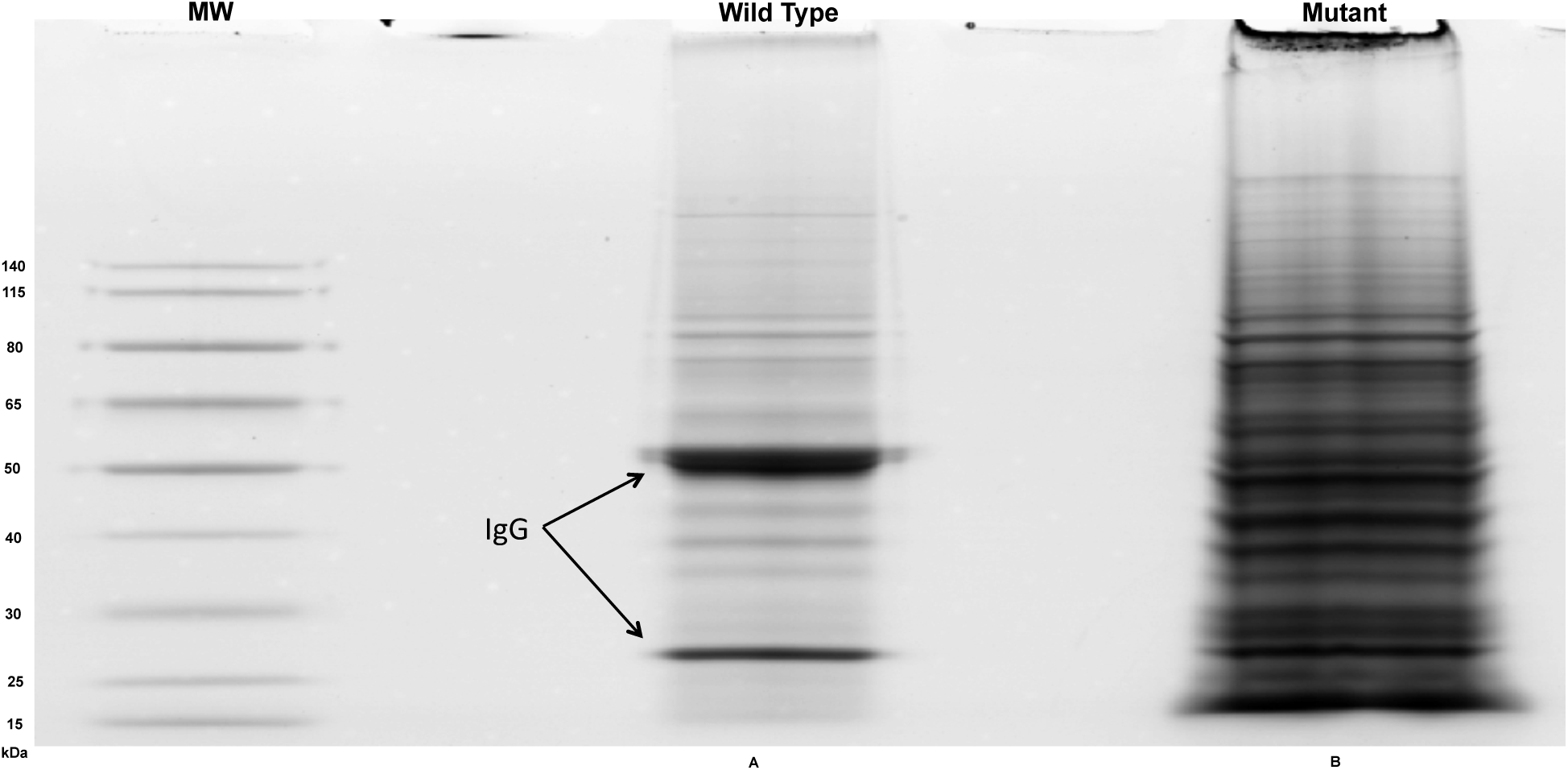
Blue silver comassie stained SDS-PAGE gel. The wells for the molecular weight markers (MW), the immunoprecipitant of wild type AKT1 expression plasmid transfected cell lysate (Wild Type) and the immunoprecipitant of mutant AKT1 expression plasmid transfected cell lysate (Mutant) are marked. The antibody (IgG) heavy and light chains are also marked. The experiment was repeated 4 times. A representative result is shown.

## 4. Discussion

The protein kinase study has to rely heavily on *in vitro* experimental protocols due to the cell impermeability of key analysis probes – the ATP analogs – and our inability to distinguish kinase reactions from one another in live cells. We set out to tackle this problem in this study. Nanoparticle-mediated intracellular delivery of A*TPγS overcomed the cell impermeability issue. Using the AKT1 protein kinase as the initial prototype, delivered A*TPγS enabled *in vivo* substrate thiophosphate tagging by mutant AKT1 with enlarged ATP binding pocket, which was created according to the Shokat chemical genetic method. Consequently, we were able to detect kinase-substrate relationship in live cells and in a kinase-specific manner.

The results pave the way to unbiased mass-spectrometry (MS) based proteomic study of kinase-substrate relationship. The AKT1 protein kinase, as discussed earlier, has a large number of substrates. Thus, as shown in Figure 8, the thiophosphate ester antibody was able to immunoprecipitate a large number of tagged proteins from the lysate of HCT116 cells expressing the mutant AKT1 protein. Comparative MS analysis of the immunoprecipitant with that of wildtype-AKT1-expressing cells will lead to unbiased identification of both known and new AKT1 substrates.

Our approach should be applicable to many other protein kinases. Due to the conservation of the ATP binding pocket across the kinome, the Shokat chemical genetic method is widely applicable. It has been applied to a large number of human protein kinases; just to name a few, EGF receptor, SRC, p38/MAPK14 and ERK1/2 (39). Sometimes, a different derivative of the ATP molecular was used. Consequently, the nanoparticle-mediated intracellular delivery of the bulky ATP analogs should be applicable in enabling *in vivo* analysis of these protein kinases as well.

The Shokat chemical genetic method has also been applied to other research areas. First, it has been used to develop more specific protein kinase inhibitors. Many kinase inhibitors act by competing with ATP for binding to the target kinases. Since the ATP binding pocket are well conserved across the kinome, such kinase inhibitors would likely target multiple kinases. This issue was alleviated by modifying the inhibitor to fit it into the enlarged binding pocket of the mutant protein kinase. This technique has been applied to generate inhibitors that have been used to probe the functions of more that 70 protein kinases (13). Thus, whether our approach can be used to improve this application of the Shokat chemical genetic method via more efficient intracellular delivery of the inbibitors becomes an worthy future endeavor.

Second, the principle of the Shokat method has been applied to study other ATP binding proteins (53). The DDX3 gene codes for a DEAD-box protein and is frequently mutated in human cancers. The DDX3 protein belongs to a large family of ATP-dependent RNA chaperones. The ATP binding pockets of the DEAD-box proteins in this family are very similar to one another, making it challenging to develop DDX3-specific inhibitors. It was shown that mutating the hydrophobic amino acid residues, which interact with ATP N6 position in the binding pocket, enabled the mutant DDX3 protein to bind to bulky ATP analogs (53). Our strategy of nanoparticle-mediated intracellular delivery should be able to ensure efficient delivery of the bulky ATP analog based mutant-DDX3-specific inhibitors.

Moreover, it is tempting to investigate whether our approach can use additional derivatives of the A*TP (N6-Benzyl-ATP) molecule for *in vivo* probing of other aspects of protein kinase function. A*TPγS derives from A*TP by substituting the γ-phosphate group with a thiophosphate group. The purpose of the thiophosphate group is to act as a handle for kinase substrate identification. However, there are other important aspects of kinase study, *e*.*g*., ATP binding and enzymatic activities. The ATP binding pocket is usually shielded from ATP binding in an inactive protein kinase, but becomes accessible upon activation of the kinase. For instance, in monomeric inactive EGF receptor (EGFR), the activation loop is in close proximity of the pocket and blocks ATP bnding. Upon ligand binding and dimerization, the C-lobe of one EGFR molecule disrupt this autoinhibitory conformation of the other EGFR molecule, enabling ATP binding and kinase activation (54). However, current ATP binding and kinase activity analyses rely on the usage of ATP analogs, and cell impermeability of these analogs renders *in vivo* analyses impossible. Using our approach for intracellular delivery of relevant A*TP derivatives should help to alleviate this cell impermeability obstacle.

Nanoparticles have been widely used for therapeutic agent delivery in biomedical researches. For instance, the LCP nanoparticle has been used to deliver gene therapy agents such as siRNA and expression vectors (55,56). It has also been used to deliver many therapeutical chemicals such as gemcitabine triphosphate (20,21), doxorubicin and paclitaxel (57). Due the similarity between A*TPγS and gemcitabine triphosphate, we choosed to use this nanoparticle for intracellular A*TPγS delivery in this study. To the best of our knowledge, this represents a new application of nanoparticles – overcoming cell impeameability of key analysis probes to enable *in vivo* execution of previously *in vitro* experimental procedures. The principle should be applicable to a wide variety of biochemical, molecular and cellular research techniques, perhaps adopting difference types of nanoparticles for different purposes. Tremoudous previously impossible research opportunities are to be opened up.

In summary, in this study, we ventured our first step towards *in vivo* protein kinase analysis by successfully combining nanoparticle-mediated probe (A*TPγS) delivery and the Shokat chemical genetic method. The former overcomed the A*TPγS cell impermeability issue, and the latter enabled differentiation of the kinase reaction of interest from those of other protein kinases. Using the AKT1 as our initial prototype, we achieved detection of kinase-substrate relationships in live cells. It is our belief that this approach can be applied to other protein kinases and other aspects of protein kinase studies. Hopefully, the principle – using nanoparticle-mediated delivery of key analysis probes to overcome cell impermeability – will be generally applicable to many other biochemical, molecular and cellular biomedical research areas.

## 5. Conclusion

Identification of the *in vivo* substrates of specific protein kinases in intact cells offers a great understanding of intracellular signaling pathways and benefits therapeutic targets for human diseases (2). Herein, we have developed a strategy to exclusively tag the direct substrates of a protein kinase of interest in intact cells, combining the chemical genetics method and the nanoparticle delivery system to uniquely transfer γ-thiophosphorylation tagging to AKT1 substates in intact cells. Specifically, we confirmed our strategy worked on previously known AKT1 substrate using immunoprecipitation and Western blot assays. Finally, this *in vivo* substrate identification strategy can identify both known and novel substrates, offers a novel route to identify substrates of various kinases, and can be applied to the signaling cascade study.

## 5. Acknowledgments

This work was supported by National Institutes of Health (NIH) grants R15GM122006 to DW and U54 CA198999 to LH.

